# Dynamic Predictive Coding with Hypernetworks

**DOI:** 10.1101/2021.02.22.432194

**Authors:** Linxing Preston Jiang, Dimitrios C. Gklezakos, Rajesh P. N. Rao

## Abstract

The original predictive coding model of Rao & Ballard [1] focused on spatial prediction to explain spatial receptive fields and contextual effects in the visual cortex. Here, we introduce a new dynamic predictive coding model that achieves spatiotemporal prediction of complex natural image sequences using time-varying transition matrices. We overcome the limitations of static linear transition models (as in, e.g., Kalman filters) using a hypernetwork to adjust the transition matrix dynamically for every time step, allowing the model to predict using a time-varying mixture of possible transition dynamics. We developed a single level model with recurrent modulation of transition weights by a hypernetwork and a two-level hierarchical model with top-down modulation based on a hypernetwork. At each time step, the model predicts the next input and estimates a sparse neural code by minimizing prediction error. When exposed to natural movies, the model learned localized, oriented spatial filters as well as both separable and inseparable (direction-selective) space-time receptive fields at the first level, similar to those found in the primary visual cortex (V1). Longer timescale responses and stability at the second level also emerged naturally from minimizing prediction errors for the first level dynamics. Our results suggest that the multiscale temporal response properties of cortical neurons could be the result of the cortex learning a hierarchical generative model of the visual world with higher order areas predicting the transition dynamics of lower order areas.

## 1 Introduction

Predictive processing theories of cortical function [1, 2, 3, 4, 5, 6] propose that the cortex implements a hierarchical generative model of the environment. For example, the Rao & Ballard predictive coding model [3] uses a spatial hierarchical generative model to explain extra-classical receptive field effects in terms of top-down predictions of lower-level representations and prediction error minimization. There is now a rapidly growing body of experimental data from many different areas of neuroscience suggesting prediction error based interpretations of various neurophysiological phenomena, ranging from sensory processing in the retina [7] and lateral geniculate nucleus (LGN) [8] to interactions between sensory and motor cortical areas [9, 10]. There is also empirical evidence for top-down predictive signals in the cortex [11, 12, 13]. Recordings in layer 2/3 (L2/3) appear to support Rao and Ballard’s original interpretation of L2/3 neurons as error-detecting neurons [3] and top-down & bottom-up signal comparators [14]. Beyond the cortex, the idea of computing errors between top-down predictions and lower-level inputs is consistent with theories of the cerebellum [15] and models of dopamine responses as reward prediction errors [16].

Considerably less attention has been placed on explaining the temporal representational hierarchy in the cortex. Experimental evidence suggests that cortical representations exhibit a hierarchy of timescales from lower-order to higher-order areas, both across sensory and cognitive regions [17, 18, 19]. Higher order areas typically have a slower response decay with longer temporal windows [19], showing more stability in neural activities compared to lower order areas. We ask the question: could such phenomena be explained by a spatiotemporal predictive coding model? Previous models for spatiotemporal predictions were restricted to signal processing in retina and LGN [7, 8]. Other models such as sparse coding [20, 21] and independent component analysis (ICA) [22] have been shown to produce oriented space-time receptive fields from natural image sequences, but these models collapse an entire image sequence to a single vector input and do not model the temporal dynamics between images; therefore, these models cannot make predictions into the future given a single input. A previous spatiotemporal predictive coding model based on Kalman filtering [23] did incorporate state transitions but the model was not hierarchical and since it was trained on limited data, it is unclear if such a model can generalize well to unseen data.

Recent advances in deep learning [24] have spurred several efforts to learn spatiotemporal hierarchies from sensory data. Lotter et al. [25] developed a deep learning model called “PredNet” for learning a hierarchical predictive coding-inspired model of natural videos. After training, the model was shown to produce a wide range of visual cortical phenomena and motion illusions. However, in PredNet, higher level neurons predict lower level *prediction errors* rather than neural activities, making it unclear what the underlying generative model is. It is also unclear if PredNet learns a slow-fast temporal response hierarchy as found in the cortex. A different model, proposed by Singer et al. [26] and later extended into hierarchies [27], is trained by making higher layers predict lower layer activities: after training, model neurons in different layers displayed different levels of tuning properties and direction selectivity similar to neurons in the dorsal visual pathway. Similar to the sparse coding and ICA models discussed above for spatiotemporal sequences, the Singer et al. model also uses a sequence of images as a single input to the network, and the hierarchy of timescales is hard-coded (higher level neurons predict future lower level neural activities by receiving a fixed-length chunk of neural activities as input).

Here, we address the limitations of previous models by introducing a new **dynamic predictive coding model** with two innovations: (1) The model uses a hierarchical spatiotemporal generative model in which higher level variables generate **entire sequences** of lower level variables by predicting their transition function. This is analogous to hierarchical hidden Markov models (HHMMs) [28, 29] where a single higher level latent variable generates a sub-HHMM that has its own transition dynamics (and can potentially generate a “sub-sub-” HHMM recursively). Thus, higher level neurons in our model’s temporal hierarchy predict the **dynamics** of lower levels rather than directly predicting lower level activities; (2) A single transition function parameterization (e.g., a single transition matrix) is too rigid and cannot adapt to different linear and nonlinear dynamics. We therefore leverage recent work on “HyperNetworks” [30] in machine learning: these are neural networks that generate weights for another neural network. However, generating an entire set of high-dimensional synaptic weights may not be neurally plausible. Instead, our model uses a hypernetwork for top-down modulation of lower-level synaptic weights by predicting a low-dimensional vector of mixture weights for a group of learned transition matrices. Such an interpretation of hypernetworks is more in line with recent ideas in neuroscience on context-dependent modulation of population dynamics for task-switching [31] and motor control [32]. In our model, the top-down modulation hypernetwork learns to dynamically mix a group of transition matrices, allowing the network to adapt quickly to different visual input dynamics.

In the following sections, we first introduce a single level model that uses a recurrent hypernetwork to modulate lower level transition matrices (Section 2). We then extend this approach to a hierarchy by describing a two-level model where the representation at the second level modulates first level dynamics via a top-down modulation hypernetwork (Section 3). Our results show that for both the single level model and the hierarchical model, the network learns a spatiotemporal generative model of natural video sequences and develops space-time receptive fields similar to simple cells in the primary visual cortex, displaying both separable and inseparable (direction-selective) space-time receptive fields [33]. Furthermore, a temporal hierarchy, as exemplified by longer timescales and greater stability of second level responses compared to first level responses, emerges naturally from minimizing prediction errors for natural image sequences without any explicit smoothness assumptions, hardwired time constants or other computational constraints (Section 4). We conclude in Section 5 with the implications of our work and future directions.

## 2 Single Level Model with Recurrent Modulation

We first describe a single-level dynamic predictive coding model based on recurrent modulation of transition dynamics. The generative model in this case factorizes as follows (Figure 1(a)):

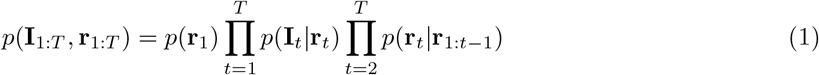

where **I**_1:*T*_ denotes the input sequence from time 1 to *T* and **r**_1:*T*_ are the latent (or hidden) state vectors, which are represented by neural activities in the predictive coding network. We use a linear spatial (or image) generative model with sparse coding [20] for *p*(**I**_*t*_|**r**_*t*_) and a recurrent hypernetwork [30] to parameterize the temporal generative model *p*(**r**_*t*_ **r**_1:*t−*1_). We describe the two parameterization choices in the following sections and conclude with the inference and learning procedure for the model based on prediction error minimization.

**Figure 1:**
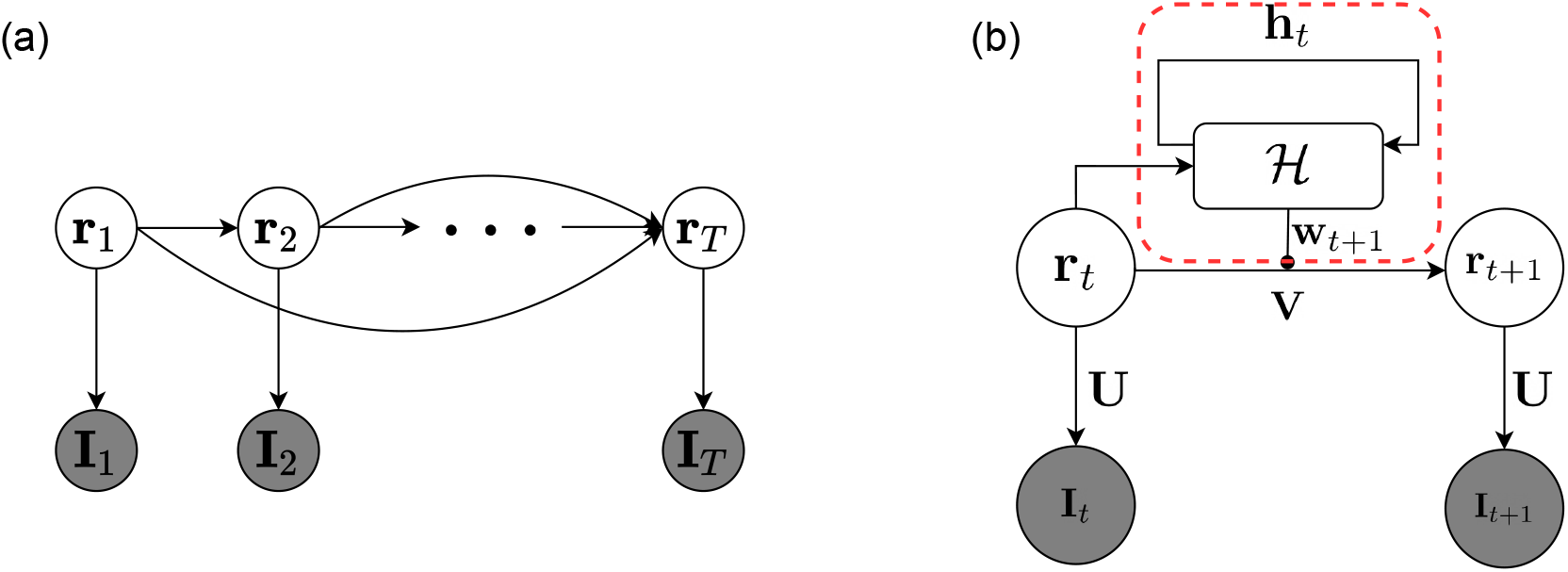
Single level predictive coding model with recurrent modulation. White nodes represent latent variables, shaded nodes represent input observations. (a) Graphical model characterizing input generation process for the single layer model; (b) Parameterization of the single layer model. Red dashed box denotes the recurrent modulatory hypernetwork that generates modulation weights **w**_*t*+1_ given the current state **r**_*t*_ and recurrent state **h**_*t*_.

### 2.1 Spatial Generative Model with Sparse Coding

Our model uses the same spatial generative model as sparse coding [20] and predictive coding [3]:

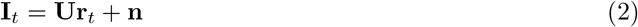

where **I**_*t*_ is the input image at time *t*, the columns of **U** represent spatial filters, **r**_*t*_ is a sparse hidden state vector, and **n** is zero mean Gaussian noise. **U** is assumed to be overcomplete, i.e. **U** has more columns than the image dimension. Inference and learning of sparse coding models minimize the following objective:

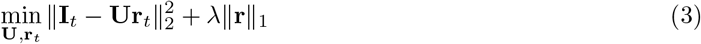

where the first term represents image prediction errors and the second term denotes the sparsity penalty on **r** weighted by a parameter *λ*. Filters learned through Equation 3 have previously been shown to closely resemble the localized, orientation-selective, bandpass receptive fields found in the mammalian primary visual cortex [20].

### 2.2 Temporal Generative Model with Recurrent Modulation

The temporal generative model consists of (1) a group of *K* transition matrices 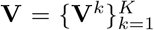 and (2) a recurrent function ℋ (implemented by a hypernetwork) that generates the linear mixing weights for these matrices (Figure 1(b) red dashed box). At each time step, ℋ takes the current state **r**_*t*_ and the recurrent state **h**_*t*_ representing past state history to generate the next set of mixture weights and the next recurrent state:

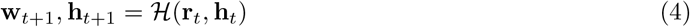

The transition matrices are then linearly weighted by **w**_*t*+1_ to produce a single transition matrix **V**_*t*+1_, which is in turn used to generate the next-step hidden state **r**_*t*+1_ given **r**_*t*_:

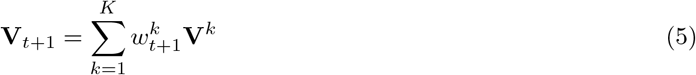

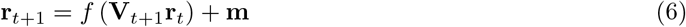

where **m** is zero mean Gaussian noise and *f* is a function (in our simulations, we use a ReLU function, which rectifies negative values to zero and leaves other values unchanged). Compared to the sparse coding model which learns a dictionary of spatial features, the model above aims to learn a dictionary of transition matrices 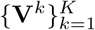 to capture the temporal dynamics of the spatial features. During inference for a given image sequence, the hypernetwork ℋ infers the optimal mixture weights for combining these transition matrices to explain the image sequence, as described below.

### 2.3 Inference & Learning as Prediction Error Minimization

Given the generative model above and the assumed parameterization, we can write down the overall optimization objective as a loss function based on prediction errors over space and time:

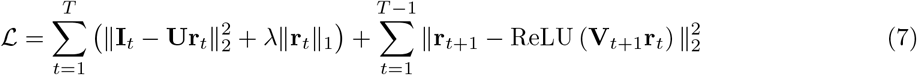

where **V**_*t*+1_ is the time-specific transition matrix generated by the modulatory hypernetwork. The squared error terms in Equation 7 correspond to the spatial and temporal prediction error respectively. At the first time step, the maximum a posteriori (MAP) estimate of **r** is inferred entirely from the spatial prediction error:

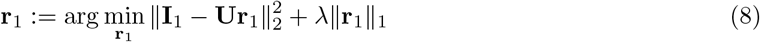

At each subsequent step *t*, we take a Bayesian filtering approach and infer the MAP estimate of **r**_*t*_ by minimizing ℒ with respect to **r**_*t*_ through gradient descent:

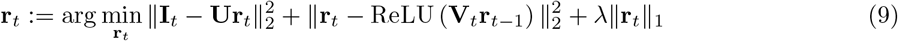

After inferring **r**_*t*_ for the whole sequence, the parameters are estimated by minimizing Equation 7 using the inferred **r**_*t*_ and performing gradient descent with respect to **U, V**, and the hypernetwork ℋ. We summarize the inference and learning procedure in Algorithm 1.

## 3 Hierarchical Model with Top Down Modulation

In this section, we extend the single level model to a two-level hierarchical model where the second level neurons influence the transition dynamics of the first level neurons via a top-down modulatory hypernetwork. We expect the second level neurons to have stable activation if the top-down predicted dynamics matches the dynamics at the first level (which in turn is determined by the input sequence). When the first level dynamics changes, we expect the resulting prediction errors to cause the second level neurons to adapt their activations and generate new transition dynamics for the lower level in order to minimize the prediction errors.

### Algorithm 1 Single Layer Model: Inference & Learning

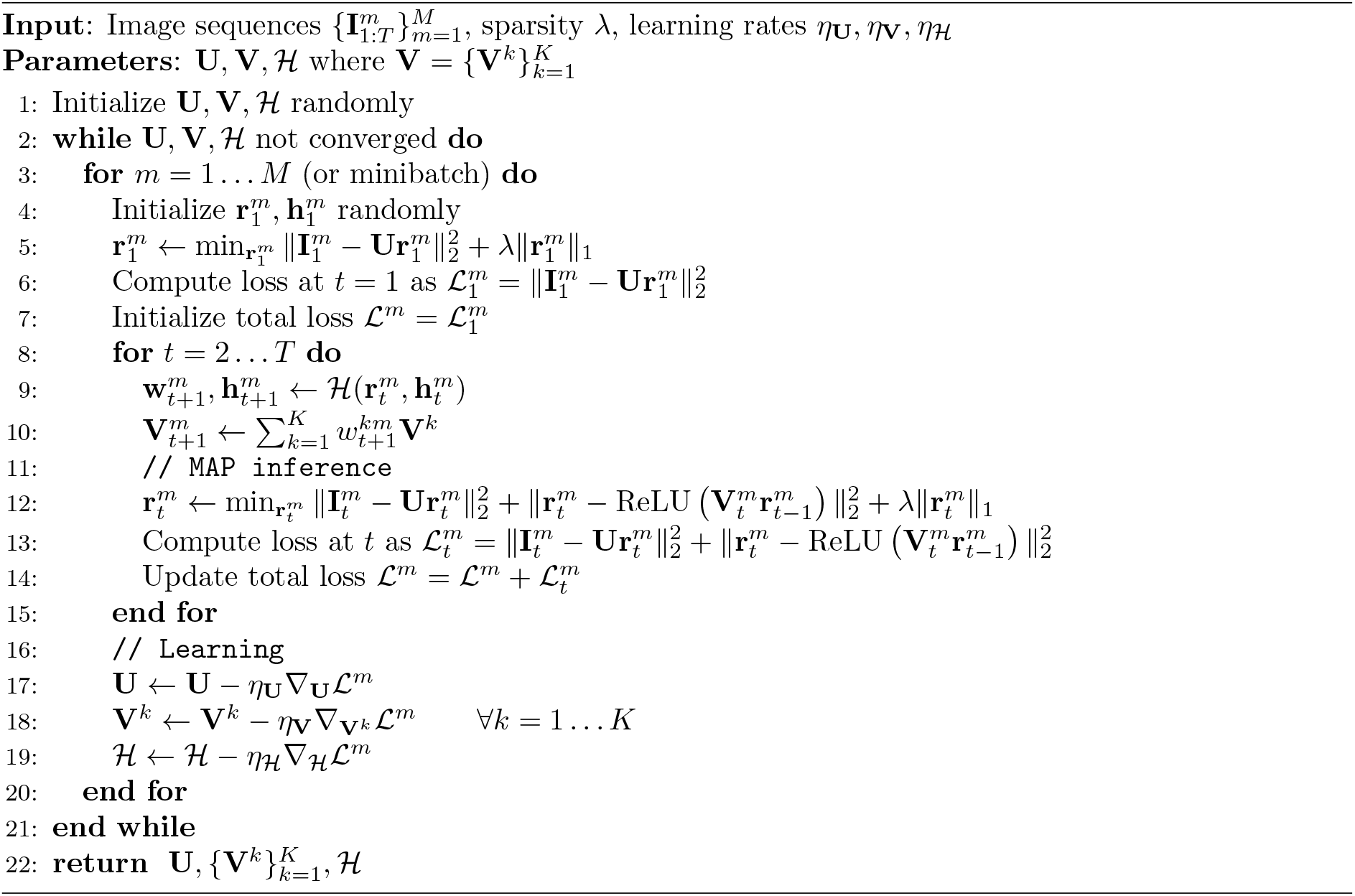

The hierarchical generative model uses a single second level hidden state variable **r**^(2)^ to influence a sequence of first level hidden state variables 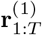 (Figure 2(a)). We assume that the two-level generative model factorizes as follows:

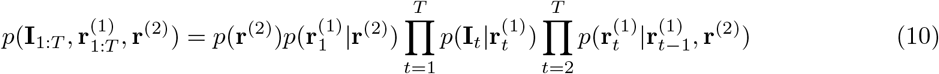

**Figure 2:**
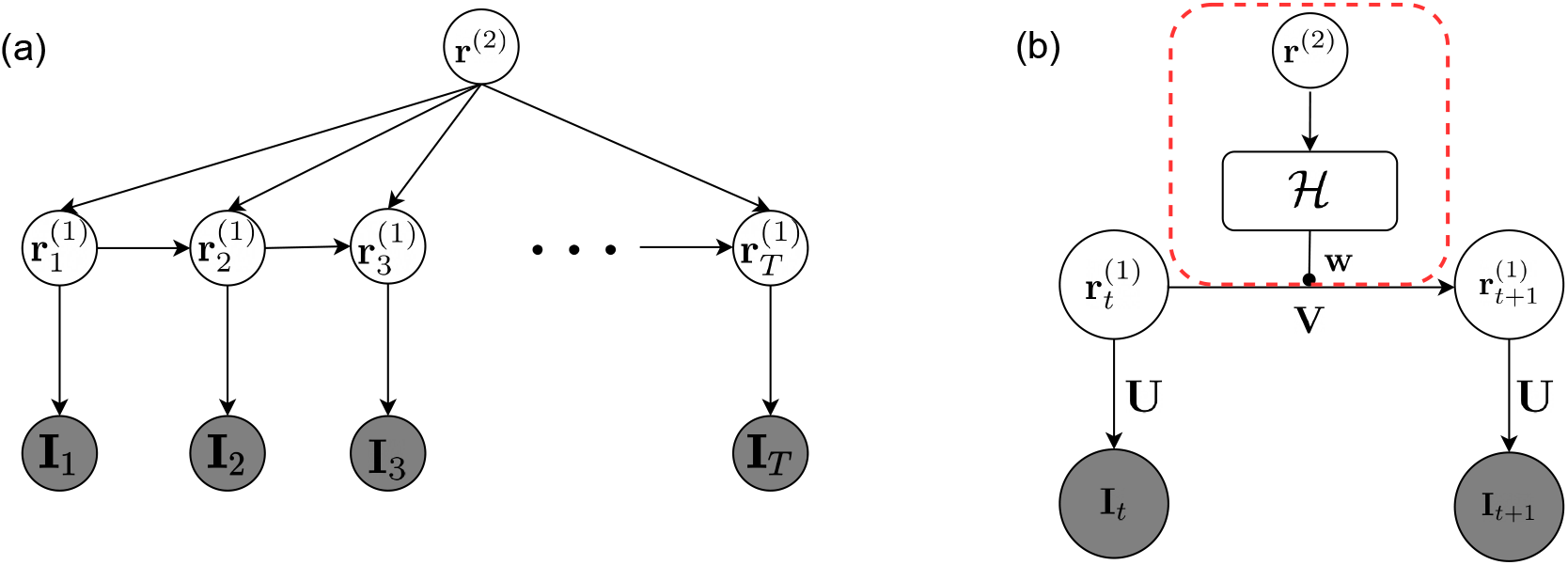
Two-level hierarchical predictive coding network. White nodes represent latent variables, shaded nodes represent observation. (a) Graphical model of the two-layer network; (b) Parameterization of the two-layer network. Red dashed box denotes the feedforward top-down modulatory hypernetwork that generates modulation weights **w** given the second level neural activities **r**^(2)^.

Note that given the second level top-down modulation, we no longer need to condition the next state 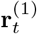 on all of the past history of states 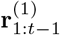 since the second level state **r**^(2)^ captures this history. Therefore, the first level state 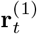 depends only on previous state 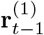 and the higher level state **r**^(2)^. For the results described in this paper, we make the further simplification that 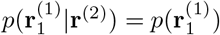.

### 3.1 State Transition Parameterization

For parameterizing 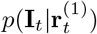, we assume the same spatial generative model (with sparse coding) as the single layer model above. For the temporal generative model, instead of the recurrent hypernet function used in the previous model, we use a non-recurrent function (implemented by a feedforward hypernetwork) that maps the second level state to the first level dynamics (Figure 2(b), red dashed box). Specifically, we parameterize the first level transition dynamics in the generative model as:

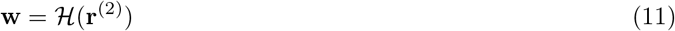

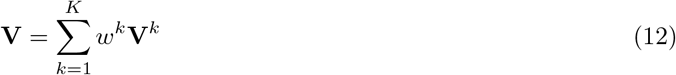

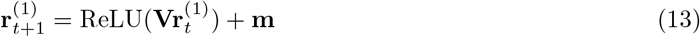

where **m** is zero mean Gaussian noise. Note that although the mixture weights **w** for the transition matrices are not time-dependent in this generative model, due to their dependence on the second level state vector **r**^(2)^, the weights produced by the hypernetwork during inference can change from one time step to the next as **r**^(2)^ is inferred from data.

### 3.2 Second Level Inference

Inference in the hierarchical model is based on minimizing the following prediction error loss function:

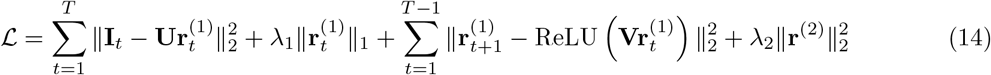

where 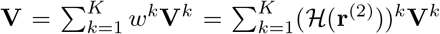 is the top-down modulated transition matrix, *λ*_1_ is the sparsity penalty parameter for 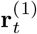 and *λ*_2_ is the Gaussian prior penalty parameter for **r**^(2)^.

Inference of 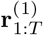 and learning of **U, V** and ℋ proceed as in Algorithm 1, after replacing the recurrent modulation of the transition matrices in Step 9 with top-down modulation: 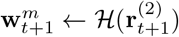. For MAP inference of **r**^(2)^, we perform gradient descent on ℒ with respect to **r**^(2)^ and update **r**^(2)^ based on the first level prediction error between 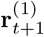 and its prediction, along with the zero mean Gaussian prior penalty on **r**^(2)^, at each time step *t*:

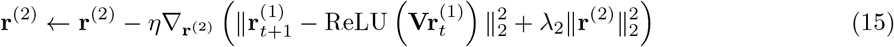

where *η* is the gradient descent rate for inference of **r**^(2)^.

Figure 3 illustrates the inference process for both layers of the hierarchical dynamic predictive coding network that implements inference for our hierarchical generative model. The network generates top-down and lateral predictions through the black arrows. If the input sequence is predicted well by the top-down hypernetwork-generated transition matrix **V**, the second level response **r**^(2)^ remains stable due to small prediction errors (Figure 3(a)). When a non-smooth transition occurs in the input sequence, the resulting large prediction errors are sent to the second level through feedforward connections (red arrows, Figure 3(b)), driving changes in **r**^(2)^ to predict new dynamics for **r**^(1)^.

**Figure 3:**
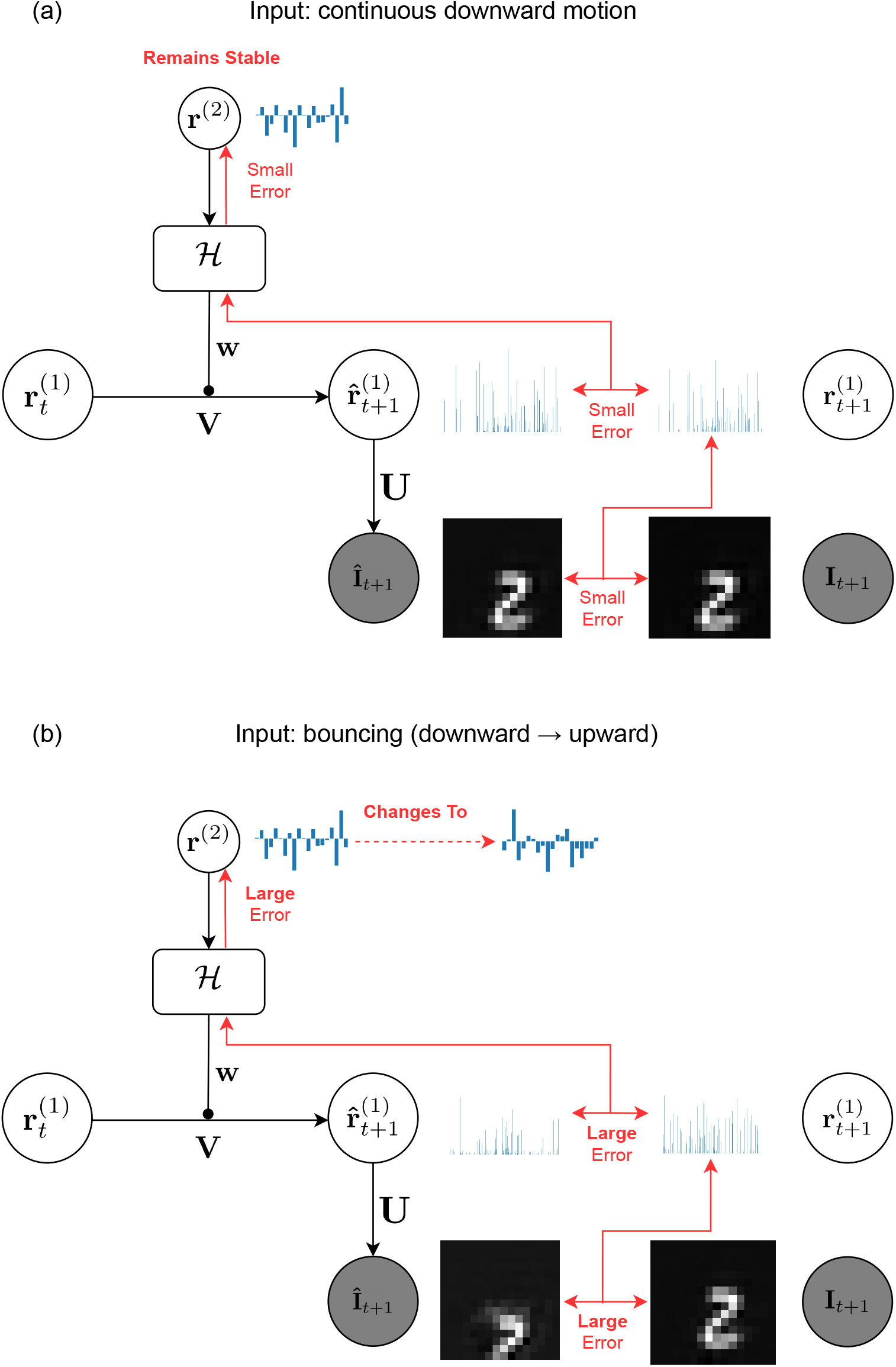
Inference in the two-level dynamic predictive coding model. Black arrows represent connections conveying top-down and lateral predictions. Red arrows represent feedforward connections conveying prediction errors. At each time step *t*, the second level **r**^(2)^ generates first level transition dynamics through the hypernetwork ℋ. The first level predicts the next state 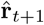 based on the top-down predicted dynamics and the current state 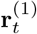. The predicted state 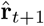 in turn generates an input prediction **Î**_*t*+1_. Prediction errors are computed between the prediction **Î**_*t*+1_ and the actual input **I**_*t*+1_ and sent through the feedforward connections to update the first and second level neural activities to 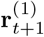 and 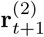 respectively. (a) Prediction error is small when the input sequence is accurately predicted by transition dynamics predicted by the second level. As a result, **r**^(2)^ remains stable. (b) When the dynamics of the input sequence changes, the resulting large prediction error at the first level is conveyed to the second level where it drives changes in **r**^(2)^ to minimize the error.

Our hierarchical network extends to continuous state spaces the framework of hierarchical Hidden Markov models (HHMMs) which have previously been defined for discrete state spaces [28, 29]. The hypernetwork can be seen as generating a continuous-valued HMM for the lower level based on a single higher level continuous-valued state. The “end state” for the lower-level HMM is indicated by large prediction errors at the lower level, signaling a need for a state change at the higher level in order to minimize prediction errors.

## 4 Results

We trained both the single layer model and the hierarchical model on 5,000 natural video segments extracted from a video recorded by a person walking on a forest trail (source: YouTube https://youtu.be/oSmUI3m2kLk; frame size: 10 × 10 pixels; segment length: 20 frames (∼ 0.7 seconds)). The video frames were spatially and temporally whitened to simulate retinal and LGN processing following the methods in Olshsausen & Field [20] and Dong & Atick [8]. For testing, we reserved 1,500 video segments that were non-overlapping in space and time with the training video segments.

We used *K* = 5 “basis” transition matrices 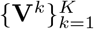 for both the single layer and the hierarchical model. We did not see significant changes in the results by increasing *K*. The size of the first level state vector **r**^(1)^ was five times the input image dimension (i.e., five-times overcomplete basis for sparse coding). For the single layer recurrent model, the hypernetwork consisted of a standard recurrent neural network (RNN) and a feedforward multilayer perceptron (MLP) that maps the hidden recurrent activities to the modulation weights **w**. Each hidden layer used a ReLU nonlinearity followed by a batch normalization layer [34] (except the last output layer). For the two-level hierarchical model, the hypernetwork was a single MLP with the exact same architecture as the one in the one-layer model (with the second level state vector as the input). Additional details on model parameterization and training are provided in Appendix A.

### 4.1 Dynamic Transition Matrix Generates Better Long Term Predictions of Natural Videos

We first tested our hypothesis that allowing the transition matrix to be dynamic (time-varying) instead using a static transition matrix as in traditional Kalman filtering (Figure 4(a)) boosts prediction performance. We used the single-level model with recurrent hypernetwork modulation as described in Section 2 and trained the model on our natural videos dataset. We computed the mean squared error (MSE) for next time-step image prediction, defined as:

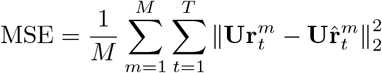

where *M* is the number of sequences. Figure 4(b) shows that the dynamic model outperforms the static model in pixel-wise prediction in both the training and the test sets. The dynamic model does have an advantage in that it uses more parameters than the static model, but note that the size of the transition matrix is pre-determined by the size of the state vector, which is the same for both models.

**Figure 4:**
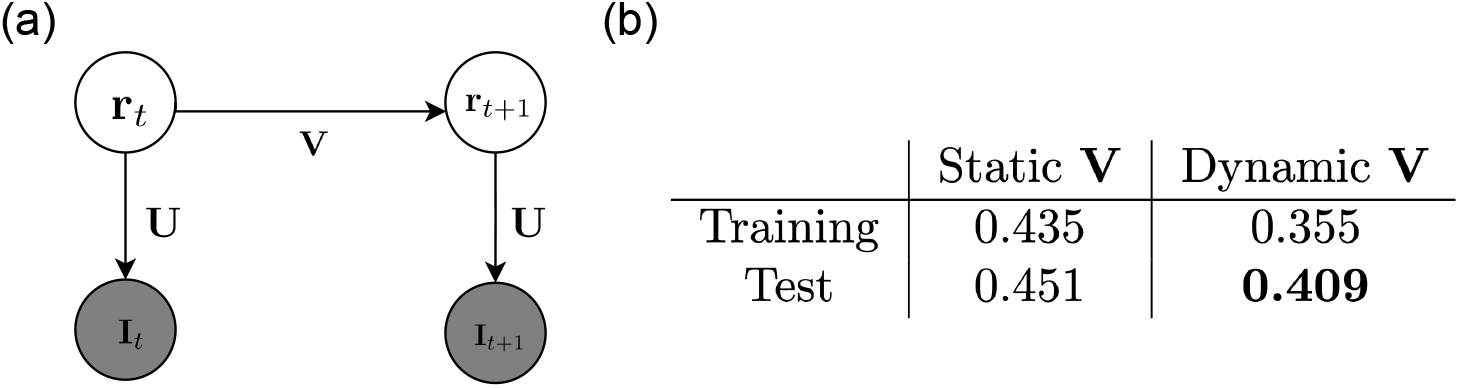
Predictive coding models with static vs. dynamic transition matrices. (a) Parameterization for a standard Kalman filtering model. The transition matrix **V** is typically assumed be static (after learning). It has the same dimensions as any of the transition matrices **V**^*k*^ in the dynamic transition model. (b) Comparison of mean squared prediction error (see text) on the training and test sets. The dynamic model outperforms the static model on both sets.

The difference in performance between the two models is more noticeable in long sequence predictions without any inputs to correct the state estimates. In this case, prediction performance depends on whether the model has inferred the correct dynamics of the input sequence in the first few input frames.

We plotted two example sequences in Figure 5 for qualitative comparison: the input sequences were turned off halfway after 10 steps (red dashed line). For the latter half of the sequence, we set **r**_*t*_ = 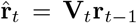 with no input to correct prediction errors. For both sequences, the dynamic transition model inferred the correct input transition dynamics (rightward for Sequence 1, upward and leftward for Sequence 2). On the other hand, though the prediction results are similar across both models in the first half when inputs are available, the static model either predicted a wrong direction (Sequence 1) or predicted no motion and blurry images (Sequence 2) in the second half when no inputs are available. These results suggest that a dynamic transition matrix generated by a hypernetwork for each time step provides greater flexibility and accuracy in capturing lower level dynamics compared to a learned but static transition matrix.

**Figure 5:**
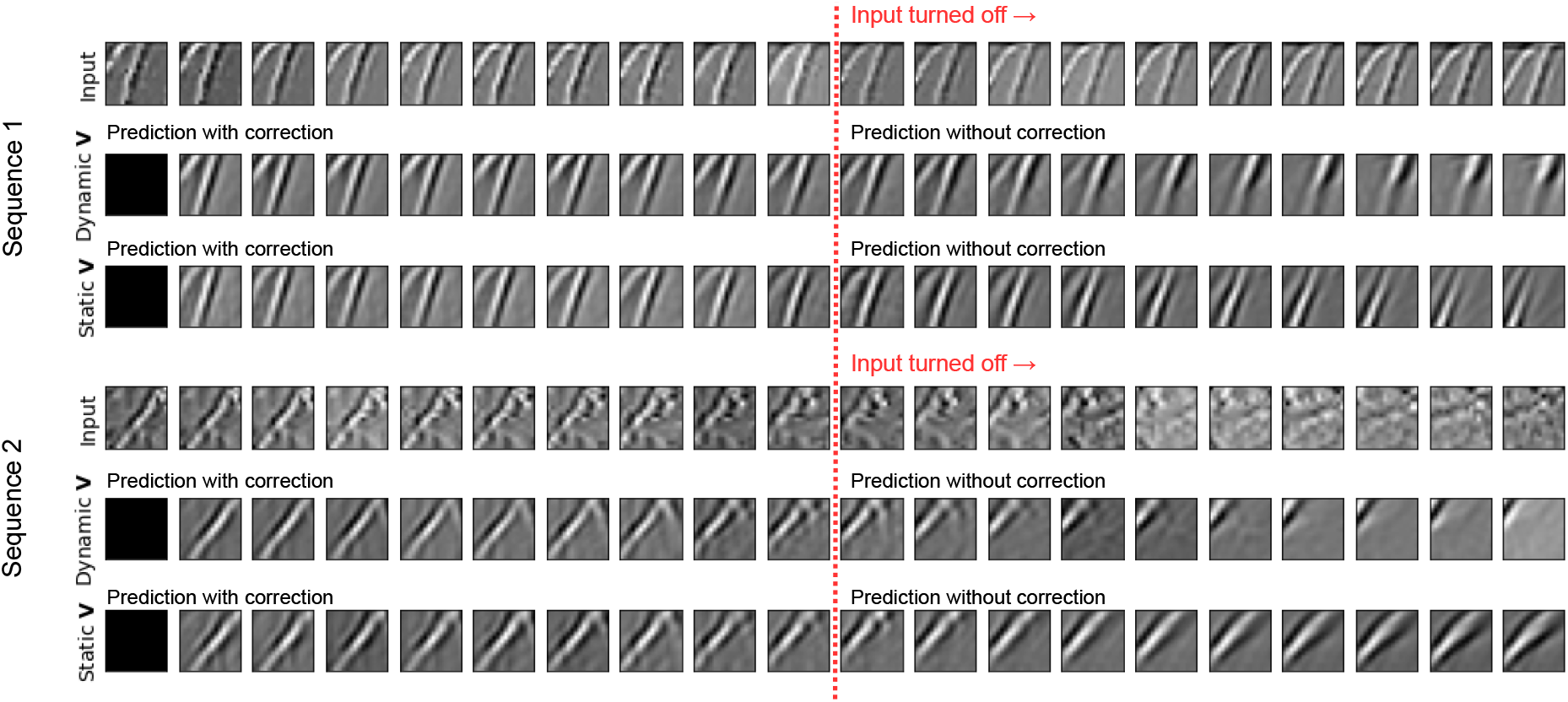
Comparison of Predictive Coding Models with Dynamic versus Static Transition Matrices. Prediction performance on two sequences of patches extracted from our test natural video dataset are shown. For each sequence: first row: actual input sequence; second row: prediction of the input by a single level model with a dynamic transition matrix; third row: prediction of the input by a single level model with a static transition matrix. Red dashed line denotes the moment when the input was turned off, after which the state vector can no longer be corrected using input prediction errors. For both sequences, the dynamic transition model correctly predicts input dynamics after input is turned off. The static transition model either predicted incorrect dynamics (top) or predicted static blurry images (bottom).

### 4.2 V1 Simple-Cell-like Space-Time Receptive Fields in First Level Model Neurons

We investigated whether the space-time receptive fields (STRFs) of the first level neurons in the dynamic model learned from natural videos resemble the STRFs of simple cells in primary visual cortex (V1). To measure the STRFs of the first level neurons representing **r**^(1)^, we ran a reverse correlation experiment [35, 36] with a continuous 30-minute natural video clip extracted from the same forest trail natural video dataset used for training but not shown to the model during either training or testing. We ran the experiments with both the single level model (Section 2) and the hierarchical model (Section 3).

Figure 6 shows examples of STRFs learned by the single level model with a recurrent hypernetwork (part (a)) and the hierarchical model with a non-recurrent hypernetwork (part (b)). For comparison, Figure 6(c) shows the STRFs in the primary visual cortex (V1) of a cat (adapted from a figure by DeAngelis et al. [33]). DeAngelis et al. categorized the receptive fields of simple cells in V1 to be space-time separable (Figure 6(c) left) and inseparable (Figure 6(c) right). Space-time separable receptive fields maintain the spatial form of bright/dark-excitatory regions over time but switch their polarization: the space-time receptive field can thus be obtained by multiplying separate spatial and temporal receptive fields. Space-time inseparable receptive fields on the other hand exhibit bright/dark-excitatory regions that shift gradually over time, showing an orientation in the space-time domain. Neurons with space-time inseparable receptive fields are direction selective.

**Figure 6:**
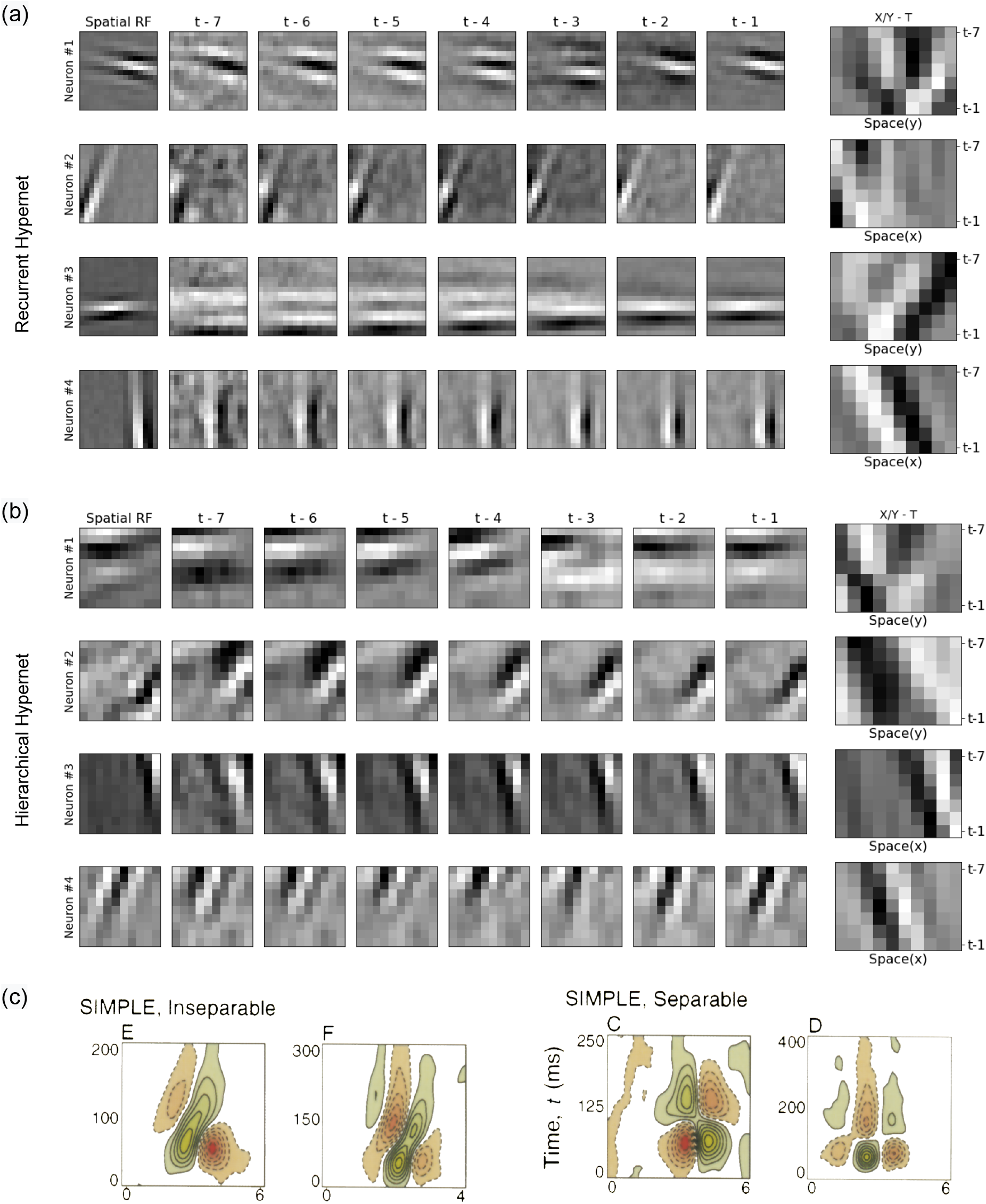
Examples of space-time receptive fields (STRFs) learned by first level neurons in the dynamic predictive coding model. (a) Example STRFs of first level neurons in the single level dynamic predictive coding model with a recurrent hypernetwork; (b) Example STRFs of first level neurons in the hierarchical dynamic predictive coding model; (c) Space-time plots of receptive fields of four simple cells in the cat primary visual cortex (adapted from [33]). (a) and (b): First column: spatial RFs of selected model neurons (each spatial RF is a column of **U**, reshaped as 10 × 10 image). Next 7 columns: spatiotemporal receptive field of the same model neurons (images show averaged stimuli occurring up to ∼ 250 milliseconds prior to current response). Last column: Space-time receptive fields of the same model neurons obtained by collapsing either the *X* or *Y* dimension (see text for details).

The model neurons in the first level of our dynamic predictive coding models learned similar V1-like separable and inseparable STRFs as revealed by reverse correlation mapping. Figures 6(a) and (b) show the STRFs learned by the single level and hierarchical model respectively. These STRFs were computed by weighting input frames for the seven time steps before the current time step by the current response 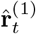 and averaging these weighted frames: the resulting averaged spatiotemporal receptive field is shown as a 7-image sequence labeled *t* − 7, *t* − 6, …, *t* − 1 in (a) and (b) (∼ 250 milliseconds in total). The first image labeled “Spatial RF” shows the spatial receptive field for each of these model neurons. To identify separability and inseparability, we took the spatiotemporal *X Y T* receptive field cubes and collapsed either the *X* or *Y* dimension, depending on which axis had time-invariant responses. The last column in Figures 6(a) and (b) shows the *X/Y − T* receptive fields of these model neurons. The dynamic predictive coding model learns both separable (rows 1, 2 part(a); row 1 part(b)) and inseparable (rows 3, 4 part(a); rows 2, 3, 4 part(b)) receptive fields. Additional STRFs of model neurons are provided in Appendices B and C.

### 4.3 Longer Timescale Responses and Generative Capabilities of Second Level Model Neurons

To derive our hierarchical model for dynamic predictive coding, we used a generative model in which a second level state generates an entire lower level sequence of states through the top-down modulatory hypernetwork. In this generative model, the higher level state remains stable during the transitions of lower level states. When the higher level state changes, a new lower level sequence is generated with different transition dynamics. We hypothesized that the different timescales of neural responses observed in the cortex [17, 18, 19] could be an emergent property of the cortex learning a similar hierarchical generative model.

We tested our hypothesis in our trained hierarchical model. Figure 7(a) shows an example natural video sequence from the test set and the hierarchical model’s responses at each level for each time step. As seen in the figure, the first level activities **r**^(1)^ change rapidly as the natural image stimulus moves in the video sequence. The second level activities **r**^(2)^ on the other hand remain stable after the initial adaptation to stimulus motion. Since the input sequence in this case continued to follow the same dynamics, the transition matrix predicted by the second level activities **r**^(2)^ continues to be accurate for the input sequence, leading to small prediction errors and little or no change in **r**^(2)^. Note that we did not place any smoothness constraints or enforce longer time constants for **r**^(2)^ - the longer timescale response and stability are entirely a result of the higher level learning to encode lower level sequences and predicting lower level transition matrices (lower level dynamics).

**Figure 7:**
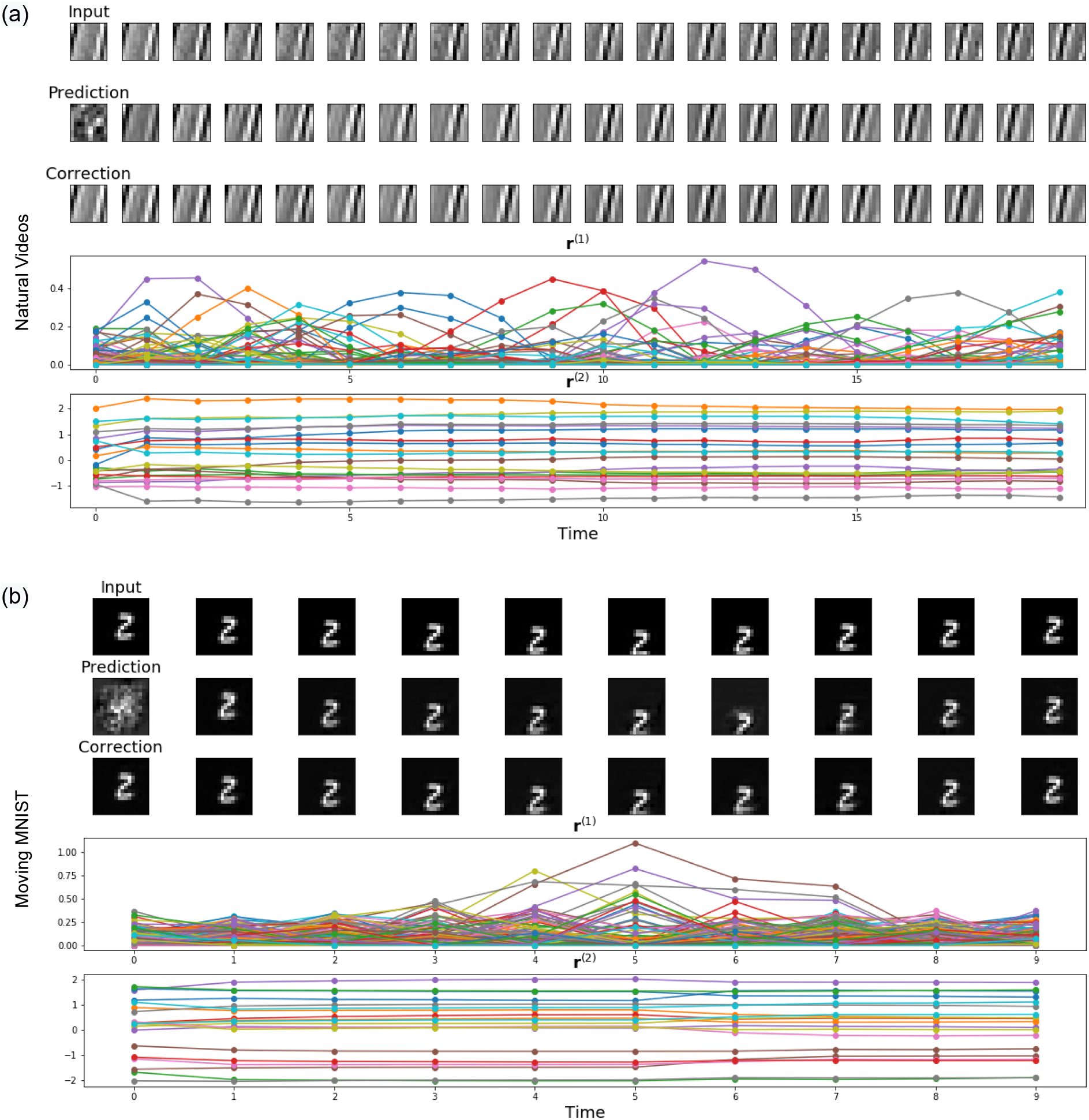
Different timescales of responses at first and second level of the hierarchical model. (a) Results from a natural video sequence. (b) Results from a Moving MNIST sequence. For both parts (a) and (b): First row: input sequence; second row: hierarchical model’s prediction; third row: generated image after error correction; fourth row: first level model neuron activities **r**^(1)^. Each plotted line represents the activity of a separate model neuron over 10 time steps; fifth row: second level model neuron activities **r**^(2)^. For both image sequences, the second level activities capture longer timescale phenomena and are consequently much more stable than the first level activities. Changes in second level activities only occur when there is a need to adapt to a new type of sequence (e.g., the digit ‘2’ moving up instead of down) in order to minimize prediction errors.

To more clearly illustrate how prediction errors can drive changes in second level model neurons to generate new lower level dynamics, we trained our hierarchical model on artificial sequences of handwritten digits from the Moving MNIST dataset [37]. The video sequences consisted of 10 16 × 16 image frames per sequence. Each frame only contained a single digit and the digit identity did not change within the sequence. We trained a hierarchical model with 400 first level neurons; all other parameters remained the same as before.

Figure 7(b) shows the trained hierarchical model’s responses to an example Moving MNIST sequence from the test set. Similar to the responses for the natural video sequence in Figure 7(a), the first level activities show fast changes while the second level shows longer timescale responses and greater stability. At time *t* = 5, the input sequence changes as the digit “bounces” against the lower boundary and starts to move upwards. Notice how the second level predicts a downward motion at *t* = 6 (prediction row, Figure 7(b)), resulting in a large prediction error as the input digit had already begun the upward motion. The gradient descent inference process for the second level causes **r**^(2)^ to change to minimize the prediction error. For the rest of the time steps, **r**^(2)^ remains stable and correctly generates the transition matrix for upward motion of the digit. These examples suggest that the progressively longer time scales and greater stability of neural responses as one ascends the visual cortical hierarchy could be the result of the cortex learning a hierarchical spatiotemporal generative model where higher level neurons encode entire sequences of lower level neural activities. Changes in neural responses in higher order areas occur when the current higher level responses can no longer predict lower level dynamics accurately.

We also investigated the generative capabilities of the hierarchical model by sampling different second level representations **r**^(2)^ and using the resulting transition matrices (obtained via the hypernetwork) to generate sequences of 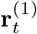 at the first level, starting from a fixed initial state 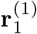. We sampled 10 different second level **r**^(2)^ from the prior 𝒩 (0, 1). We then estimated the first level representation 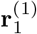 for a natural image patch **I**_1_ (same as the first step in Figure 7(a)). The hypernetwork generated 10 different transition matrices from the 10 sampled **r**^(2)^ which were used at the lower level to generate 10 sequences of 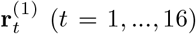 and the corresponding image sequences. Figure 8 shows the image sequences. Each row is a different sampled image sequence generated by a different **r**^(2)^ (first column) and the corresponding modulation weights **w** = ℋ(**r**^(2)^) (second column). No inputs were provided to the model besides the image to initialize 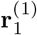. The image sequences generated by the different second level representations **r**^(2)^ (which were held fixed throughout the sequence) show a variety of transformations related to form, motion, and speed. These results indicate that the hierarchical network learned to encode different transition dynamics using different”embedding” vectors **r**^(2)^ at the second level.

**Figure 8:**
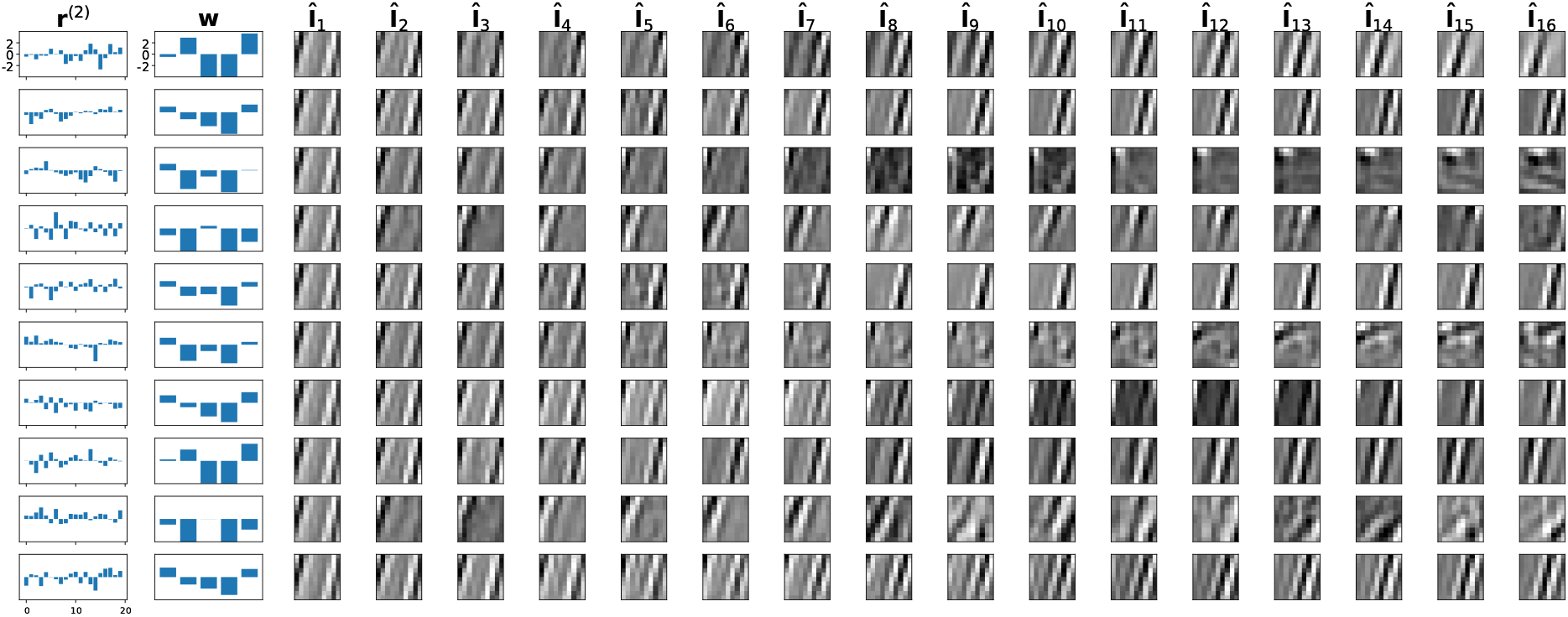
Generative capabilities of the second level representation r^(2)^. First column of plots: sampled **r**^(2)^; second column: the predicted transition dynamics, shown as the mixture weights **w** for the transition matrices 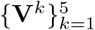. Image sequences: Each row of images starts with the same initial image generated by a fixed first level activation vector 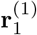. The next 15 images in each row are generated by applying that row’s transition matrix (generated by that row’s **r**^(2)^) to the first level activation vector 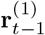 from the previous time step and predicting an image (via **U**) from the resulting activation vector 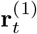. Note that a variety of transformations related to form, motion, and speed are generated by the different sampled second level representations.

## 5 Discussion

This paper presents a new dynamic predictive coding model for learning spatiotemporal input sequences. The model allows dynamically modulated temporal transitions by using a hypernetwork to generate transition matrices. By adopting the objective of minimizing spatiotemporal prediction errors (equivalent to MAP inference), the model learns a hierarchical representation encoding spatiotemporal sequences at different timescales: lower level model neurons adapt quickly to account for rapidly changing input sequences while activities at the second level remain more stable and predict entire first level sequences. Our predictive coding model can be seen as a neural implementation of a continuous-state hierarchical hidden Markov model (HHMM), with hypernetworks allowing a single higher level state to generate the transition matrix for an entire lower level sequence (sub-HHMM). During inference, lower levels of the model follow the top-down predicted dynamics and implicitly reach an “end state” [28, 29] when the input strongly disagrees with the prediction of the input, resulting in a large prediction error. The feedforward pathways in the network convey this prediction error to higher levels which correct their neural activities and generate new dynamics for the lower levels to minimize prediction error.

When the dynamic predictive coding model was exposed to thousands of natural video sequences, the first level of the model developed V1 simple cell-like properties including both separable and inseparable space-time receptive fields [33]. The longer timescale and greater stability of neural responses in higher order cortical areas [17, 18, 19], often cited as evidence for a hierarchy of timescales in the brain, emerge naturally in our network as the higher level model neurons learn to encode sequences of lower level activities by minimizing prediction errors of the lower level dynamics. After learning, sampling the higher level neural activities generates distinct lower level sequences.

Together, these results show that the dynamic predictive coding model can successfully learn a hierarchical generative model for natural videos based on a temporal hierarchy of representations. The use of hypernetworks in our model to modulate lower level dynamics suggests a new interpretation of neuromodulation of cortical synapses, a topic worthy of further study. Other directions for future research include a more quantitative comparison of the space-time receptive fields learned by the predictive coding model neurons and those found in the visual cortex, and exploring a deeper hierarchical model to explain response properties of higher order cortical areas using the principle of prediction error minimization.

## Appendix A. Network Architecture and Model Details

### Single Level Recurrent Hypernet-Based Model Architecture

Below is a summary of the dimensionality of the input and hidden variables in the single layer model used in our experiments:

- Input image **I**_*t*_ ∈ ℝ^100^.
- Spatial filters matrix **U** ∈ ℝ^100×500^, state vector **r**_*t*_ ∈ ℝ^500^.
- Transition matrices 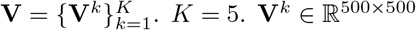..

The hypernetwork consisted of a standard RNN with 400 hidden neurons (with ReLU activation) and a feedforward multilayer perceptron (MLP) with 4 hidden layers. The input layer of the MLP had 400 neurons (taking the recurrent hidden state as the input) and the output layer had 5 neurons (generating the mixture weights in R^*K*^). All intermediate hidden layers of the MLP had 100 neurons (with bias terms and ReLU nonliearity, followed by batch normalization [34]). Changing the number of hidden layers in the MLP did not significantly affect the results.

### Hierarchical Top-down Hypernet-Based Model Architecture

Below is a summary of the dimensionality of the input and hidden variables in the two-level hierarchical model used in our experiments:

- Input image **I**_*t*_ ∈ ℝ ^100^.
- Spatial filters matrix **U** ∈ ℝ ^100×500^, first level state vector 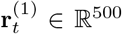, second level state vector **r**^(2)^ ∈ ℝ ^20^.
- Transition matrices 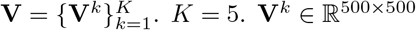.

The hypernetwork was a feedforward MLP that had the same dimensions as the MLP in the single layer model above, except that the input layer had 20 neurons (the input is the second level state vector). As in the single level model, changing the number of layers in the MLP did not significantly affect the results.

### Training Details

We used stochastic gradient descent for both learning the spatial filters matrix **U** and inferring all latent variables (gradient descent update rate: 0.005 for **U** and **r**^(1)^, and 1.0 for **r**^(2)^). We used the Adam optimizer [38] for faster convergence of the learning process for **V** and the hypernetwork ℋ, with learning rate 0.0005 and default *β* terms (*β*_1_ = 0.9, *β*_2_ = 0.999) for both. We used *λ*_1_ = 0.0005 for the sparse coding penalty on 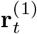 and *λ*_2_ = 0.0001 for the Gaussian prior penalty on **r**^(2)^ (see Equation 14 in text). We did not perform extensive hyperparameter tuning to optimize performance. The model is robust to other hyperparameter changes if the sparse coding shrinkage is tuned carefully. Both models (single level and hierarchical) converged after 200 epochs of iterating through the training set.

## Appendix B. STRFs in the Single Level Model

**Figure 9:**
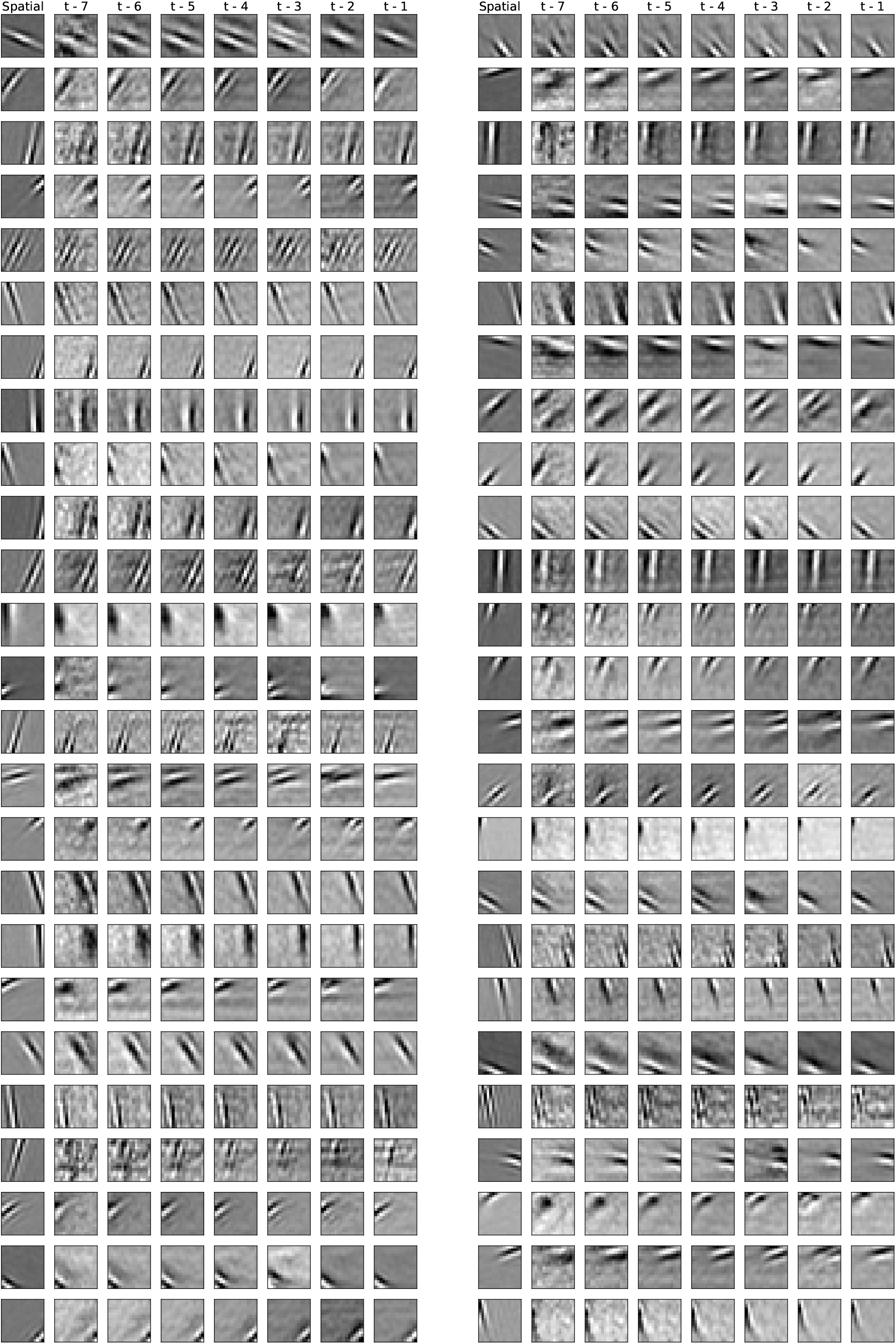
Space-time receptive fields of 50 neurons in the single level model based on the recurrent hypernetwork.

## Appendix C. STRFs in the Hierarchical Model

**Figure 10:**
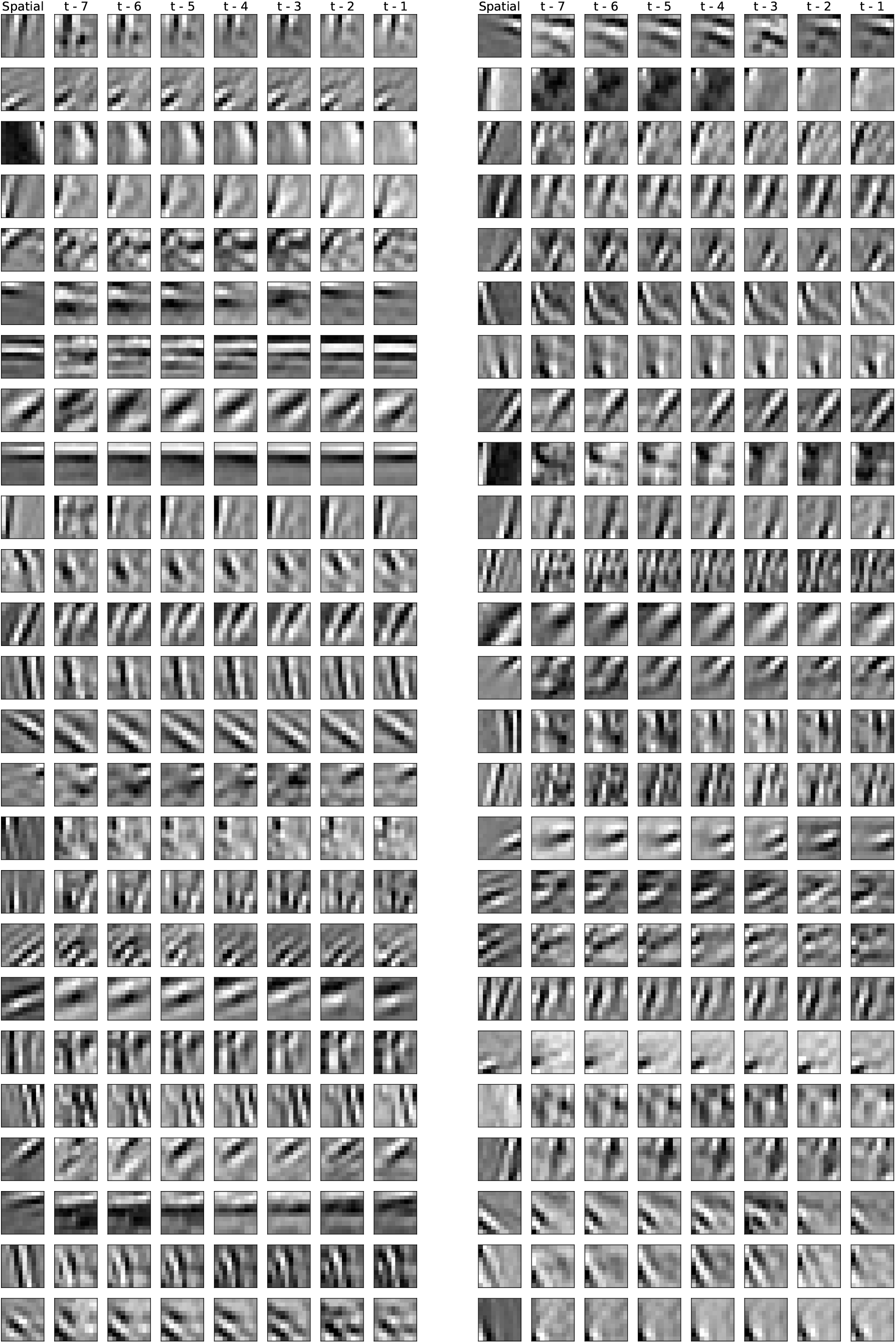
Space-time receptive fields of 50 r^(1)^ neurons in the two-level hierarchical model based on the top-down hypernetwork.

